# Antimicrobial Resistance and Virulence-Associated Genes of *Pasteurella multocida* and *Mannheimia haemolytica* Isolated from Polish Dairy Calves with Symptoms of Bovine Respiratory Disease

**DOI:** 10.1101/2024.09.25.614927

**Authors:** Agnieszka Lachowicz-Wolak, Aleksandra Chmielina, Iwona Przychodniak, Magdalena Karwańska, Magdalena Siedlecka, Małgorzata Klimowicz-Bodys, Kamil Dyba, Krzysztof Rypuła

## Abstract

**Background:** Bovine respiratory disease causes significant economic losses in cattle farming due to mortality, treatment costs, and reduced productivity. It involves viral and bacterial infections, with *Pasteurella multocida* (*P. multocida*) and *Mannheimia haemolytica* (*M. haemolytica*) as key bacterial pathogens. These bacteria contribute to severe pneumonia and are often found together. A total of 70 bacterial strains were analysed: 48 *P. multocida* and 22 *M. haemolytica*, collected from deep nasal swabs or lung and bronchial swabs from affected calves. The bacterial species were confirmed molecularly using PCR, which was also employed to detect antimicrobial resistance and virulence-associated genes. Antimicrobial susceptibility was determined using the broth microdilution method.

**Results:** Antimicrobial resistance varied between the two bacterial species studied. The highest resistance in *P. multocida* was observed to chlortetracycline 79.2% and oxytetracycline 81.3%, while *M. haemolytica* showed 63.6% resistance to penicillin and tilmicosin. Multidrug resistance among *P. multocida* was 27.1%, while among *M. haemolytica* it reached 40.9%. The most commonly observed phenotypic resistance patterns were ‘chlortetracycline, oxytetracycline’ in 37.5% of *P. multocida* and ‘ceftiofur, chlortetracycline, oxytetracycline, penicillin, tilmicosin, tulathromycin’ in 18.2% of *M. haemolytica*. The highest susceptibility was found for fluoroquinolones: *P. multocida* demonstrated 91.7% susceptibility to enrofloxacin, while 77.3% of *M. haemolytica* were susceptible to both enrofloxacin and danofloxacin. Multidrug resistance was detected in 31.4% of all tested strains. MIC_50_ and MIC_90_ determinations were performed for all tested antimicrobials. All *M. haemolytica* contained the *lkt, gs60*, and *gcp* genes. *P. multocida* carried the *sodA* gene, while the *hgbB* and *ompH* genes were present in 37.5% and 20.8% of strains, respectively. The *tetH* and *tetR* genes were observed only in *P. multocida*, at frequencies of 20.8% and 16.7%, respectively. Both species carried the *mphE* and *msrE* genes, though at lower frequencies between 6.3% and 14.6%.

**Conclusions:** This study expands the knowledge of the pathogenicity and antimicrobial resistance of *P. multocida* and *M. haemolytica* in dairy calves. *P. multocida* exhibited the highest resistance to tetracyclines, *M. haemolytica* demonstrated the greatest nonsusceptibility to penicillin. Both bacterial species were found to be susceptible to fluoroquinolones. One third of strains showed multidrug resistance.

## Introduction

Bovine respiratory disease (BRD) is a multifactorial disease involving viral and bacterial infections, that can be further aggravated by environmental factors and stress [1–5]. This leads to significant economic losses in the cattle industry due to the high costs of treatment, reduced weight gain, delayed time to first oestrus and conception, resulting in later lactation and lower milk production in cows that have recovered from BRD [6–8]. The etiology of BRD encompasses a range of viral and bacterial pathogens [9–14].

*Pasteurella multocida* (*P. multocida*) and *Mannheimia haemolytica* (*M. haemolytica*) are two of the most frequently identified bacterial pathogens in calves suffering from BRD [10, 15, 16]. These bacteria are often found as coinfections and are generally considered opportunistic pathogens [10, 17]. *M. haemolytica*, in particular, is linked to BRD outbreaks due to its production of virulence factors, such as leukotoxin, which contribute to clinical symptoms and cause severe pneumonia [18–20].

BRD is among the most common causes of antimicrobial use in dairy cattle [21–23]. According to the European Medicines Agency (EMA), Poland has one of the highest levels of antimicrobial use in food-producing animals among European Union countries [24]. The very frequent administration of these drugs and the irrational use of antimicrobials by veterinarians who do not conduct susceptibility testing contribute to increasing resistance, creating a vicious cycle. Recent research from Poland revealed that, over the course of three months on large dairy farms, individual calves received antimicrobial treatment up to two times [22]. The rising antimicrobial resistance is a pressing issue, impacting not only the health of calves but also the health of human, particularly given that *P. multocida* has zoonotic potential [25–29].

To date, there is a notable absence of published data regarding the antimicrobial susceptibility of *P. multocida* and *M. haemolytica* isolated from Polish calves suffering from BRD. This gap in the literature underscores the urgent need for detailed investigations to better understand the resistance profiles of these pathogens.

The primary aim of this study was to determine the antimicrobial susceptibility of *P. multocida* and *M. haemolytica* strains isolated from dairy calves with BRD. An additional objective was to assess the frequency of virulence-associated genes among these strains.

## Materials and methods

### Sample collection

The study included samples collected between February 2021 and January 2024. The material consisted of either deep nasal swabs or lung and bronchi swabs from calves that exhibited symptoms of BRD, such as nasal and ocular discharge, fever, cough, increased respiratory rate, enhanced lung field murmur, and respiratory wheezing. Samples from the lower respiratory tract were collected postmortem from calves that had died while showing symptoms of BRD. Swabs were preserved in Amies transport medium with charcoal (Deltalab, S.L., Rubi, Spain) and transported within 48 hours to the “Epi-Vet” veterinary diagnostic laboratory at Wroclaw University of Life Sciences.

### Isolation and initial identification

Swabs were initially cultured on Columbia Agar with 5% sheep blood (Graso, Stargard Szczeciński, Poland). The media were incubated at 37°C for 16-24 hours in an atmosphere containing 5% CO_2_. Colonies were checked for cytochrome oxidase production using the “Oxidase Test Stick” (Liofilchem, Via Scozia, Italy). The isolates were biochemically tested according to the manufacturer’s instructions using the commercial RaPID NF system (Thermo Fisher Diagnostics B.V., Landsmeer, NL). The biochemical reactivity patterns were analysed using the dedicated ERIC™ v.1.1 software (Thermo Fisher Diagnostics B.V., Landsmeer, NL). Selected isolates of *P. multocida* and *M. haemolytica* were cryopreserved in CryoBank (Mast Group Ltd., Reinfeld, Germany) and stored at -80°C for further studies. Species confirmation was subsequently performer via molecular methods.

### Genomic DNA extraction

Genomic DNA was extracted using the Genomic Mini Kit (A&A Biotechnology, Gdynia, Poland) according to the manufacturer’s instructions.

### Amplification setup

PCR amplification was performed using the Color Taq DNA Polymerase kit (Eurx, Gdańsk, Poland). The reaction mixture contained the following components: 0.625 U of Color Taq DNA Polymerase, 1 µL of template DNA, 0.1 µL of each forward and reverse primer,1 µL of dNTP mixture and 2.5 µL of 10X Buffer B (Eurx, Gdańsk, Poland). The mixture was supplemented with nuclease-free water (A&A Biotechnology, Gdynia, Poland) to a final volume of 25 µL.

### Cycling conditions

The PCRs reactions were performed in a T100TM Thermal Cycler (Bio-Rad Laboratories, Inc., Hercules, CA, USA). The PCR protocol began with an initial denaturation step at 95°C for 5 minutes. This was followed by 32 amplification cycles, each consisting of denaturation at 94°C for 30 seconds, annealing at a temperature specific to the primers used for 30 seconds and extension at 72°C for 1 minute per kilobase (kb) of the amplified product. After the completion of the cycles, a final elongation step was performed at 72°C for 10 minutes. The mixture was then cooled to 4°C for further analysis.

### Primers for gene detection

All primers used in this study were synthesized by Genomed S.A. (Warsaw, Poland). For all the genes detected, the primer pair sequences, annealing temperatures, and product sizes are listed in Table 1.

**Table 1.**
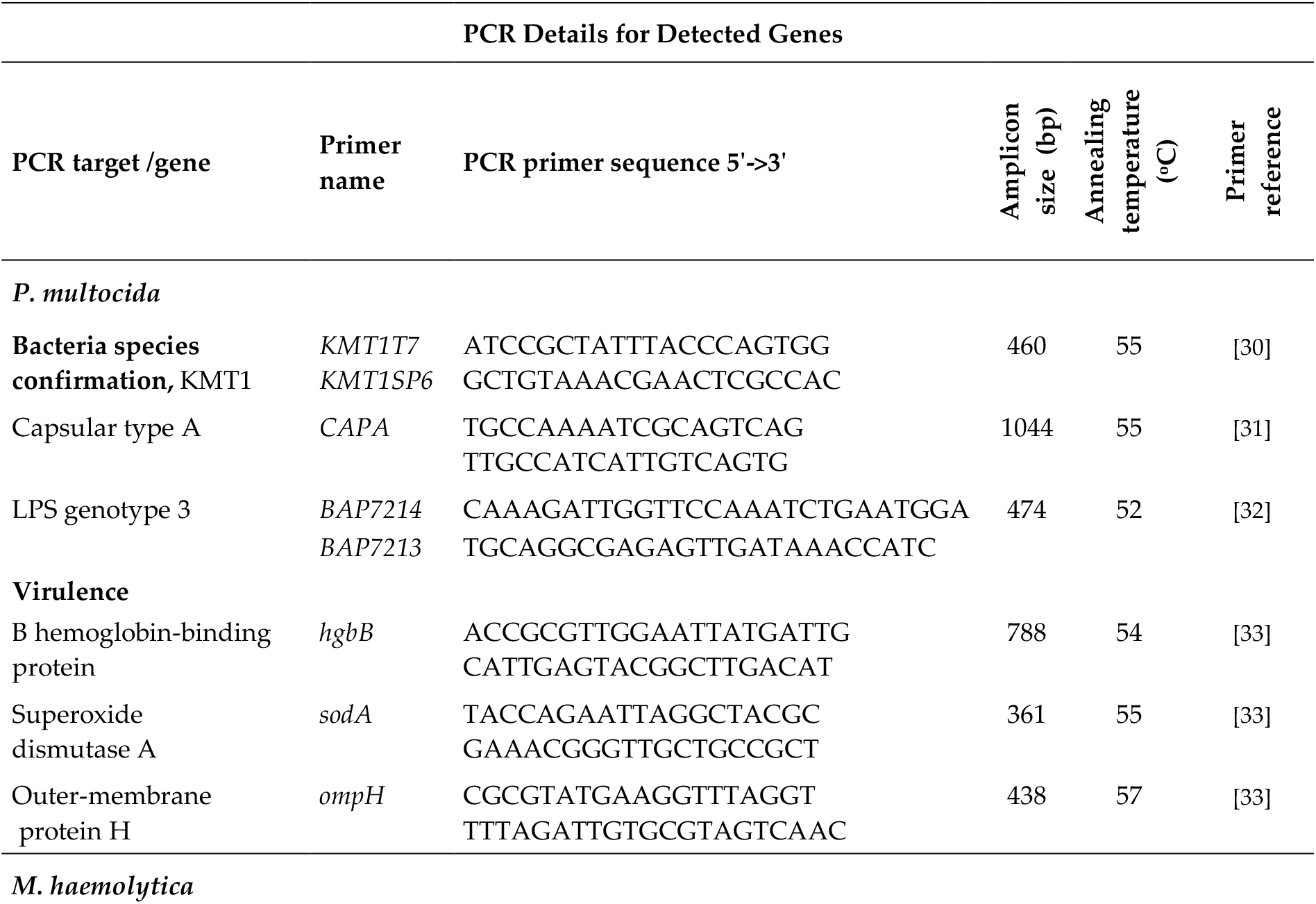
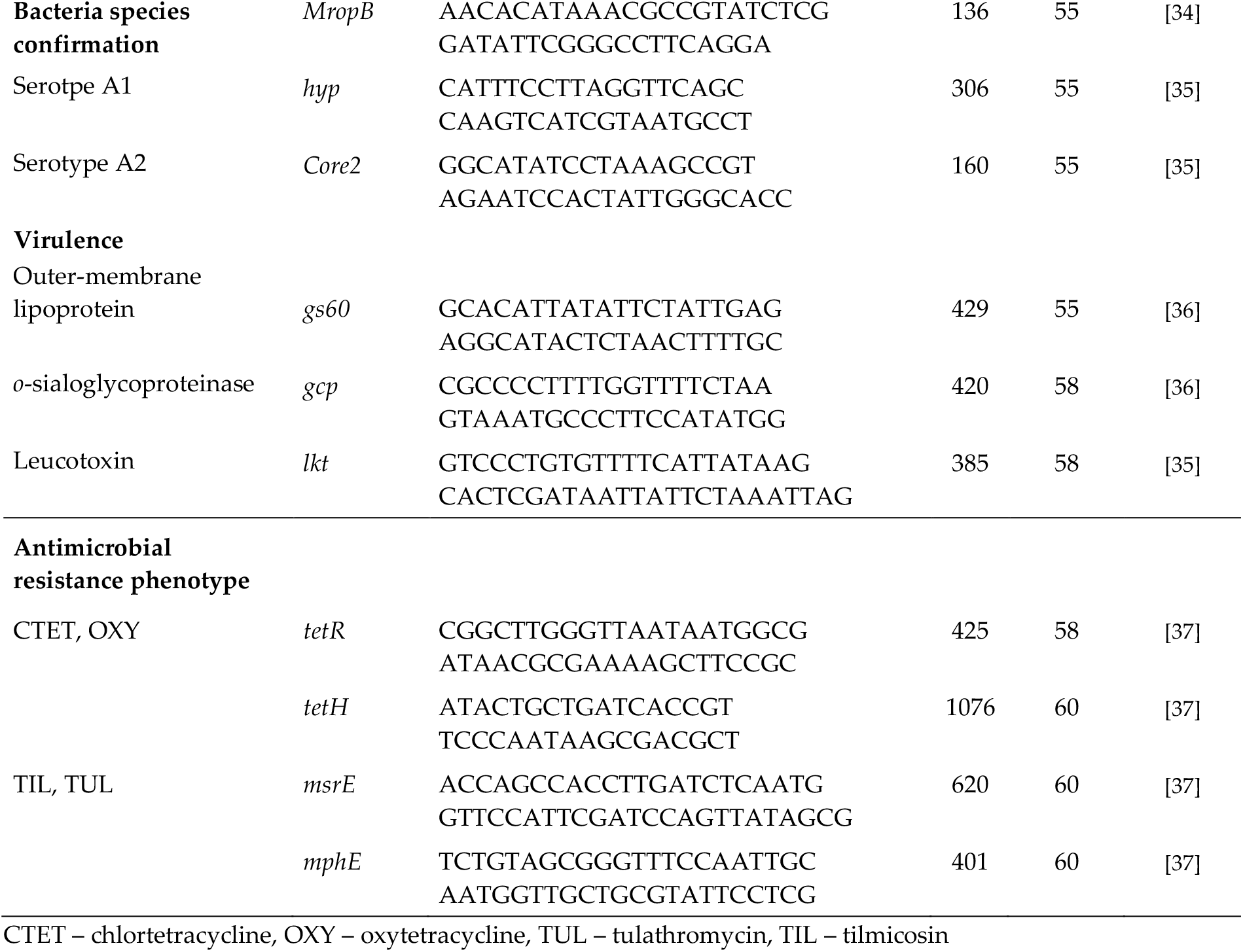
Primer sequence, amplicon size, annealing temperature, and gene accession numbers for all the genes detected in this study.

### Electrophoresis of PCR products

PCR-generated products were detected by electrophoresis in 1.5% agarose gels stained with Midori Green DNA Stain (Nippon Genetics Europe GmbH, Düren, Germany) alongside a DNA Marker 1, 100-1000 bp marker ladder (A&A Biotechnology, Gdynia, Poland) using a PowerPacTM Basic (Bio-Rad Laboratories Inc., Hercules, CA, USA) at 95 V and 400 mA. The electrophoresis results were read using the GelDoc Go Imaging System with Image Lab v.6.1 software (Bio-Rad Laboratories Inc., Hercules, CA, USA).

### Positive controls and sequencing validation

As positive controls, *P. multocida* ATCC 12945 was used for the *kmt, capA, ompH, sodA*, and *hgbB* genes, and *M. haemolytica* PCM 2685 from Polish Academy of Science was used for the *MropB, hyp, lkt, gs60* and *gcp* genes. To confirm the accuracy of the PCR results for all tested *M. haemolytica* genes, gene encoding lipopolysaccharide type 3 and resistance genes, the representative two products were selected and subsequently sent to Genomed S. A. (Warsaw, Poland) for Sanger sequencing with both forward and reverse PCR primers. The obtained sequences were analyzed using BioEdit v5.0.9 software and compared with sequences from the National Center for Biotechnology Information (NCBI) GenBank database, confirming their alignment with those available in the database.

### MIC testing methodology

The minimal inhibitory concentrations (MICs) for *M. haemolytica* and *P. multocida* were assessed using the broth microdilution method with the commercially available test plate Thermo Scientific SensititreTM BOPO6F (TREK Diagnostic Systems Ltd., East Grinstead, United Kingdom). The antimicrobials included: ceftiofur (XNL), tiamulin (TIA), chlortetracycline (CTET), gentamicin (GEN), florfenicol (FFN), oxytetracycline (OXY), penicillin (PEN), ampicillin (AMP), danofloxacin (DANO), sulphadimethoxine (SDM), neomycin (NEO), trimethoprim/sulfamethoxazole (SXT), spectinomycin (SPE), tylosin tartrate (TYLT), tulathromycin (TUL), tilmicosin (TIL), clindamycin (CLI), enrofloxacin (ENRO).

The testing procedures adhered to the guidelines established by the Clinical and Laboratory Standards Institute (CLSI) [38]. For optimal growth of the primary cryopreserved isolates, an inoculum of approximately 5 × 10^5^ cfu/mL was prepared in Thermo Scientific SensititreTM Cation Adjusted Mueller-Hinton Broth with 5% Lysed Horse Blood (Remel Inc., Lenexa, USA). The inoculated Sensititre plates were incubated in ambient air for 18–24 hours at 36 ± 1°C within a humidified chamber. Quality control is performed on the plates by the manufacturer; however, in addition, we conducted testing on a *P. multocida* ATCC 12945.

### Evaluation of MIC results

The MIC values were recorded using the SensititreTM Manual MIC Plate Reader (Thermo Fisher, Waltham, MA, USA). The results were evaluated according to the CLSI standards to assess the assessment of the susceptibility of the tested bacterial strains to the antimicrobial agents [38]. Based on these evaluations, each strain was classified as susceptible (S), intermediate (I), or resistant (R) to the respective antimicrobial. In all instances where the term “ nonsusceptible” is used, it refers collectively to both intermediate and resistant classifications. After determining the MIC values for each of the antimicrobials, population analyses were carried out to determine the MIC_50_ (MIC required to inhibit 50% of the organisms) and MIC_90_ (MIC required to inhibit 90% of the organisms) values. The multiple antimicrobial resistance (MAR) index was calculated as the quotient between the number of antimicrobials to which strains were resistant to the number of tested antimicrobials. Multidrug resistance (MDR) was defined as resistance to at least one substance in three or more antimicrobial classes.

### Statistical analysis

To assess the statistical significance of the difference in resistance to antimicrobials between the two studied bacterial species and to assess the correlation between the presence of virulence genes Pearson’s chi-square tests without Yates’ correction were performed. Similarly to check for correlations between antimicrobial resistance genes and the resistance results demonstrated in the MIC test, cross-tab analyses were performed along with Pearson’s chi-square tests without Yates’ correction (χ² test) and additionally, Pearson’s correlation. The significance level for all tests was set at α = 0.05.

Statistical analyses were performed using Statistica 13.3.721.1 (TIBCO Software Inc., Palo Alto, CA, USA). Tables and graphs were prepared using Microsoft Office Excel 2007. Figures were prepared using Adobe Illustrator and VennPainter V.1.2.0. [39].

### Ethic statement

According to Polish law (the Experiments on Animals Act from 15 January 2015, Journal of Laws of the Republic of Poland from 2015, item. 266), this study did not require the Ethics Committee’s approval. The samples used in this study were originally from cattle infection diagnostic material collected by veterinarians treating these herds.

## Results

### Tested isolates

Samples from 42 different herds were tested, all of which were collected from calves exhibiting symptoms of BRD. Three of the samples were obtained from the lower respiratory tract, while the remaining samples were deep nasal swabs. Based on phenotypic, biochemical, and molecular analyses, a total of 70 bacterial isolates were included for further analysis: 48 *P. multocida* isolates and 22 *M. haemolytica*. Based on the presence of capsule genes, all *P. multocida* isolates were assigned to serogroup A, with 93.8% (45/48) of the isolates genotypically identified as the L3 lipopolysaccharide type, while the remaining three isolates could not be classified in terms of lipopolysaccharide type. Among the *M. haemolytica* isolates, 13.6% (3/22) were classified as serotype A1, while the remaining 86.4% (19/22) were classified as serotype A2. Among the three samples from the lower respiratory tract, two strains of *P. multocida* A:L3 and one strain of *M*.*haemolytica* serotype 2 were obtained.

### Antimicrobial susceptibility

The highest susceptibility was observed for fluoroquinolones: *P. multocida* showed 91.7% (44/48) susceptibility to enrofloxacin, while 77.3% (17/22) of *M. haemolytica* strains were susceptible to both enrofloxacin and danofloxacin. The highest percentage of intermediate susceptibility to antimicrobials was observed for *M. haemolytica*, with 50.0% (11/22) of the tested strains exhibiting intermediate susceptibility to FFN and SPE. The *P. multocida* strains demonstrated the highest resistance to tetracyclines, with 79.2% (38/48) resistant to CTET and 81.3% (39/48) resistant to OXY. In contrast, the *M. haemolytica* strains showed the highest resistance to PEN and TIL, with 63.6% (14/22) resistance. The remaining susceptibility percentages are presented in Table 2.

**Table 2.** Susceptibility of the tested strains to antimicrobial agents.

| Antimicrobial | Antimicrobial Susceptibility |  |  |  |  |  |  |  |  |  |  |  |
| --- | --- | --- | --- | --- | --- | --- | --- | --- | --- | --- | --- | --- |
|  | *Pasteurella multocida* |  |  |  |  |  | *Mannheimia haemolytica* |  |  |  |  |  |
|  | S |  | I |  | R |  | S |  | I |  | R |  |
|  | n/N | % | n/N | % | n/N | % | n/N | % | n/N | % | n/N | % |
| CTET | 9/48 | 18.8% | 1/48 | 2.1% | 38/48 | 79.2% | 10/22 | 45.5% | 2/22 | 9.1% | 10/22 | 45.5% |
| OXY | 9/48 | 18.8% | 0/48 | 0.0% | 39/48 | 81.3% | 11/22 | 50.0% | 1/22 | 4.5% | 10/22 | 45.5% |
| TUL | 37/48 | 77.1% | 0/48 | 0.0% | 11/48 | 22.9% | 9/22 | 40.9% | 1/22 | 4.5% | 12/22 | 54.5% |
| TIL | 34/48 | 70.8% | 2/48 | 4.2% | 12/48 | 25.0% | 7/22 | 31.8% | 1/22 | 4.5% | 14/22 | 63.6% |
| PEN | 36/48 | 75.0% | 3/48 | 6.3% | 9/48 | 18.8% | 3/22 | 13.6% | 5/22 | 22.7% | 14/22 | 63.6% |
| ENRO | 44/48 | 91.7% | 3/48 | 6.3% | 1/48 | 2.1% | 17/22 | 77.3% | 4/22 | 18.2% | 1/22 | 4.5% |
| DANO | 42/48 | 87.5% | 4/48 | 8.3% | 2/48 | 4.2% | 17/22 | 77.3% | 2/22 | 9.1% | 3/22 | 13.6% |
| XNL | 36/48 | 75.0% | 5/48 | 10.4% | 7/48 | 14.6% | 14/22 | 63.6% | 1/22 | 4.5% | 7/22 | 31.8% |
| FFN | 38/48 | 79.2% | 9/48 | 18.8% | 1/48 | 2.1% | 9/22 | 40.9% | 11/22 | 50.0% | 2/22 | 9.1% |
| SPE | 29/48 | 60.4% | 7/48 | 14.6% | 12/48 | 25.0% | 10/22 | 45.5% | 11/22 | 50.0% | 1/22 | 4.5% |

Breakpoint values determined by the CLSI were used for the analysis. CTET – chlortetracycline, OXY – oxytetracycline, TUL – tulathromycin, TIL – tilmicosin, PEN – penicillin, ENRO – enrofloxacin, DANO – danofloxacin, XNL – ceftiofur, FFN – florfenicol, SPE – spectinomycin.

Statistically significant differences in resistance to some of the tested antimicrobial agents were observed between the bacterial species: PEN χ2 = 13.778; TUL χ2 = 9.646; OXY χ2 = 9.205; CTE χ2 = 7.956; and TIL χ2 = 6.841, with p-values < 0.05. The mentioned antimicrobials are marked with an asterisk in the following graph depicting the percentage of nonsusceptible strains (Figure 1).

**Figure 1.**
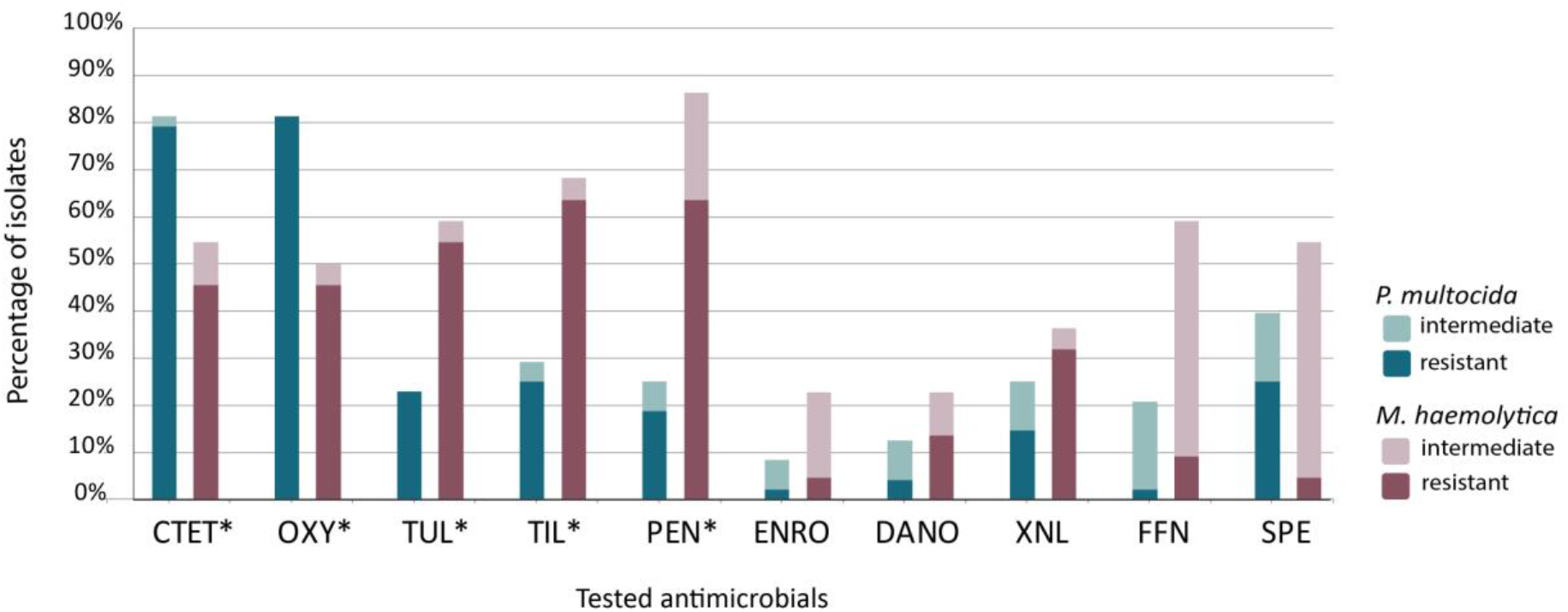
Percentages of nonsusceptibility to antimicrobials among the tested strains of *P. multocida* and *M. haemolytica*. * indicates a statistically significant difference in resistance to the given antimicrobial agent between bacterial species (p < 0.05). CTET – chlortetracycline, OXY – oxytetracycline, TUL – tulathromycin, TIL – tilmicosin, PEN – penicillin, ENRO – enrofloxacin, DANO – danofloxacin, XNL – ceftiofur, FFN – florfenicol, SPE – spectinomycin.

### MIC values

The effectiveness of individual antimicrobials was determined using MIC_50_ and MIC_90_ (Figure 2). It was not possible to identify the concentration effective at inhibiting the growth of half of the tested strains (MIC_50_) for 22.2% (4/18) of the antimicrobials tested against *P. multocida* and *M. haemolytica*, with this percentage calculated separately for each bacterial species. Similarly, determining the concentration effective at inhibiting the growth of 90% of strains (MIC_90_) was not possible for 55.6% (10/18) of antimicrobials for *P. multocida* and 61.1% (11/18) for *M. haemolytica*.

**Figure 2.**
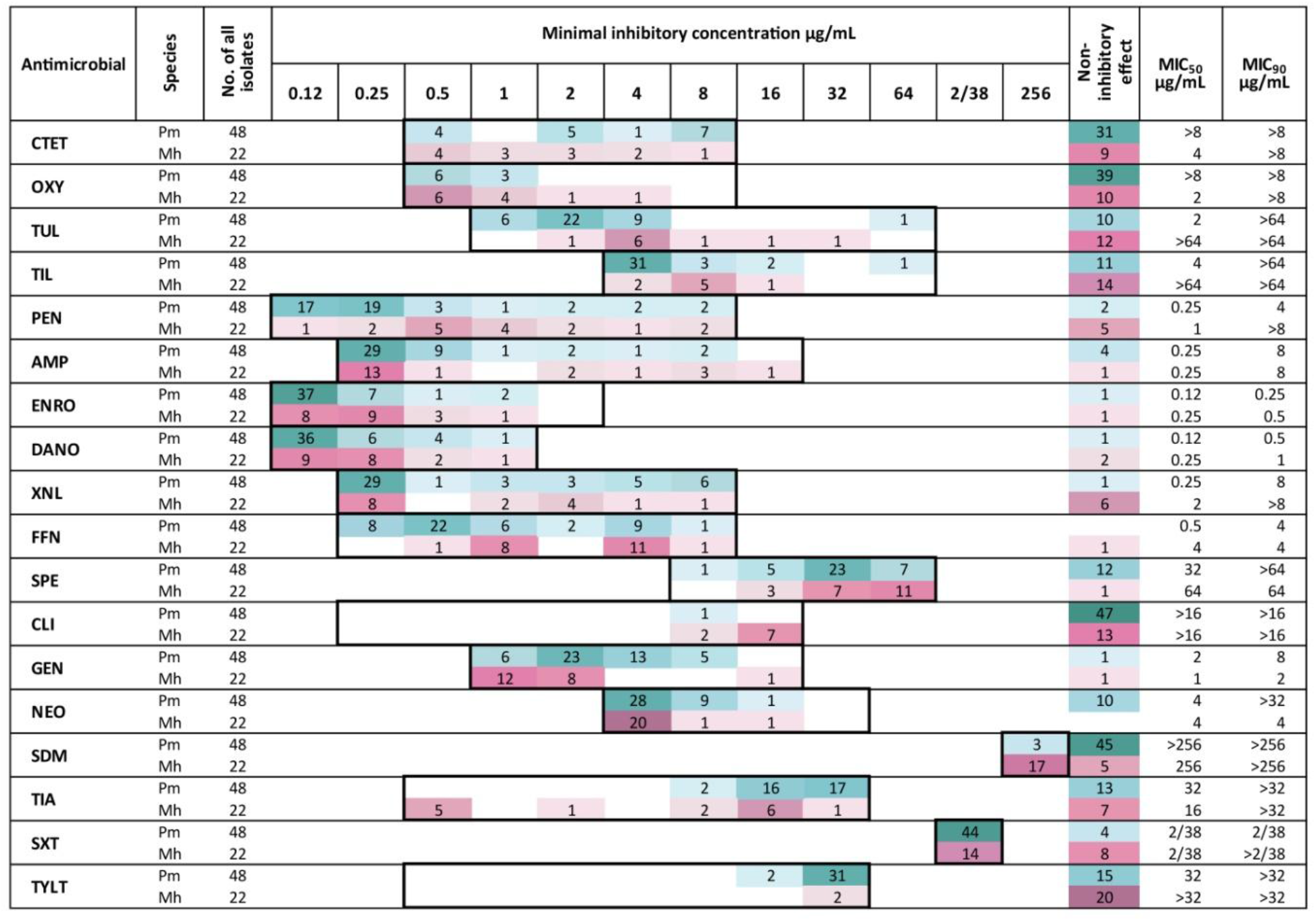
Number of bacterial strains of *P. multocida* (Pm) and *M. haemolytica* (Mh) showing growth inhibition at the specified concentrations of the tested antimicrobial agents. The number of strains not inhibited at the tested concentrations was also included. MIC_50_ and MIC_90_ values are provided. The range of tested concentrations is indicated by the boxes and the darker the color, the greater the number of strains. CTET – chlortetracycline, OXY – oxytetracycline, TUL – tulathromycin, TIL – tilmicosin, PEN – penicillin, AMP – ampicillin, ENRO – enrofloxacin, DANO – danofloxacin, XNL – ceftiofur, FFN – florfenicol, SPE – spectinomycin, CLI – clindamycin, GEN – gentamicin, NEO – neomycin, SDM – sulphadimethoxine, TIA – tiamulin, SXT – trimethoprim/sulfamethoxazole, TYLT – tylosin tartrate.

### Multidrug resistance and phenotypic resistance patterns

MDR was detected in 31.4% (22/70) of all the tested strains. Among the *P. multocida* strains, the rate was 27.1% (13/48); among the *M. haemolytica* strains, it was 40.9% (9/22). MAR index ranged from 0.275 for *P. multocida* to 0.336 for *M. haemolytica*. The most frequently noted phenotypic resistance pattern was ‘CTET, OXY,’ which was observed exclusively among *P. multocida* strains at 37.5% (18/48). Among the *M. haemolytica* strains, the most common resistance pattern was the cooccurrence of resistance to ‘XNL, CTET, OXY, PEN, TIL, TUL’, which was present in 18.2% (4/22) of the tested strains. All the recorded resistance patterns are presented in Table 3.

**Table 3.**
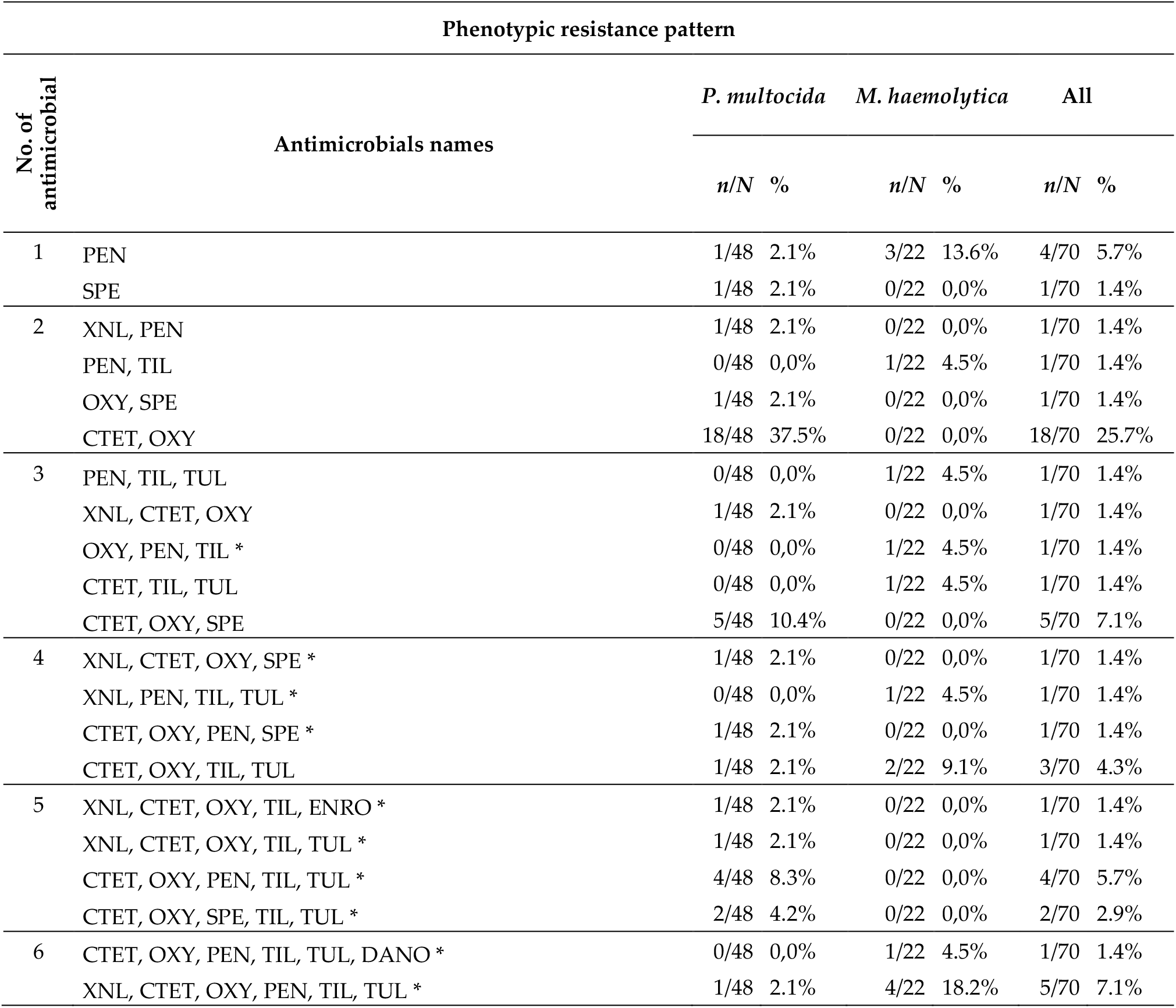
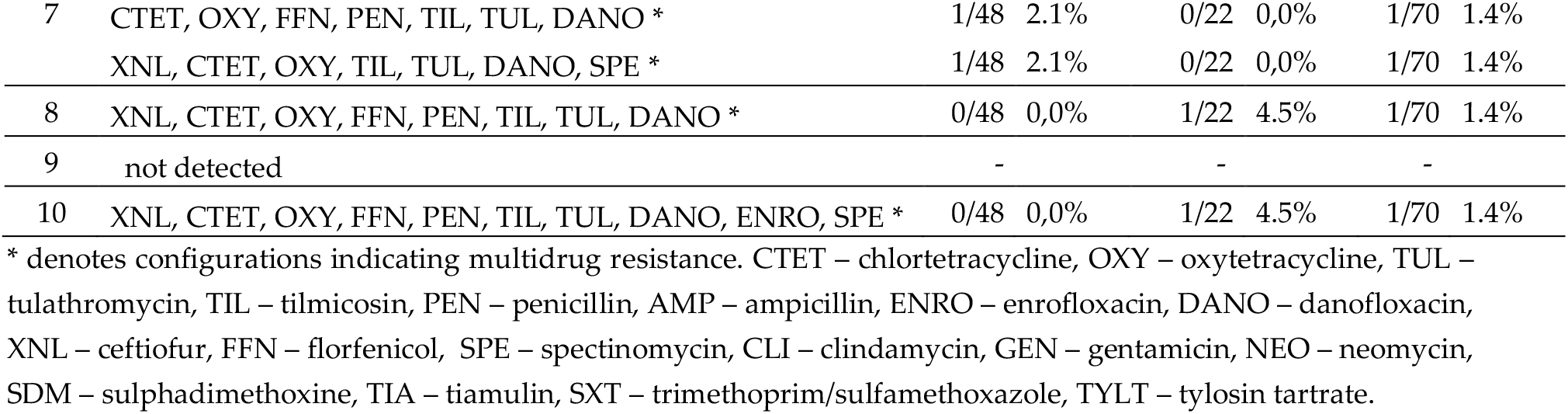
Resistance patterns detected among the tested strains of *P. multocida* and *M. haemolytica* along with their frequencies.

### Frequency of virulence-associated genes

Among the tested strains, 100% of the *P. multocida* strains possessed the *sodA* gene, while the other VAGs, namely *hgbB* and *ompH*, were found in 37.5% (18/48) and 20.8% (10/48) of the strains, respectively. All three genes together were detected in 8.3% (4/48) of the strains. The *M. haemolytica* strains contained the *lkt, gs60*, and *gcp* genes in 100% (22/22) of the cases.

### Frequency of resistance genes

The genes *tetH* and *tetR* were observed exclusively among *P. multocida* strains, at 20.8% (10/48) and 16.7% (8/48), respectively. The genes *mphE* and *msrE* were detected in *P. multocida* at 6.3% (3/48) and 14.6% (7/48), respectively, and in *M. haemolytica* at 9.1% (2/22) each.

### Associations

No statistically significant correlations were found between the pairs *hgbB* and *ompH*. Moreover, the one hundred percent presence of the remaining VAGs and the low frequency of antibiobials resistance genes rendered inferences based on the statistical correlation tests potentially unreliable. Regardless, these tests did not show any significant correlation at p < 0.05. However, the cooccurrence of the *tetR* or *tetH* genes was always consistent with phenotypic tetracycline resistance in MIC assays.

Therefore, we decided to present the results in the form of sets showing cooccurrences, as illustrated by the Venn diagrams (Figure 3, Figure 4).

**Figure 3.**
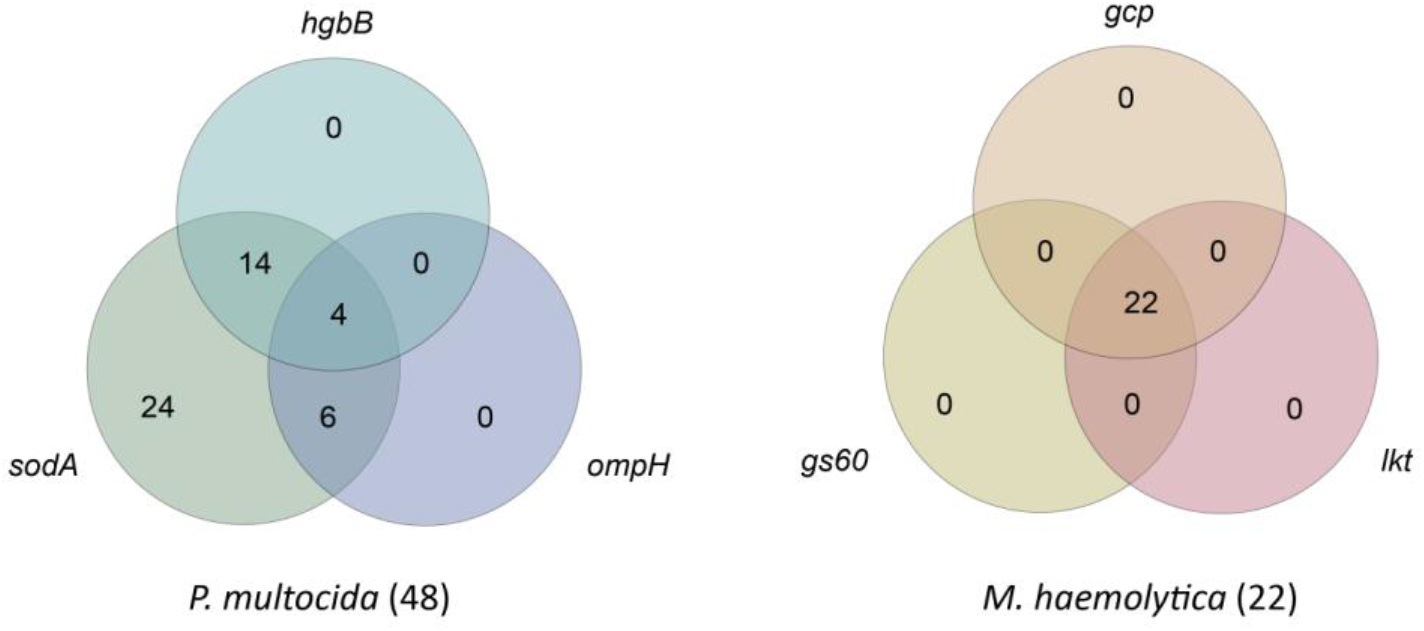
Venn diagrams showing the cooccurrence of virulence-associated genes in the studied strains of *P. multocida* and *M. haemolytica*. The common parts indicate the types of configurations, while the numbers in the respective fields represent the frequency of each configuration.

**Figure 4.**
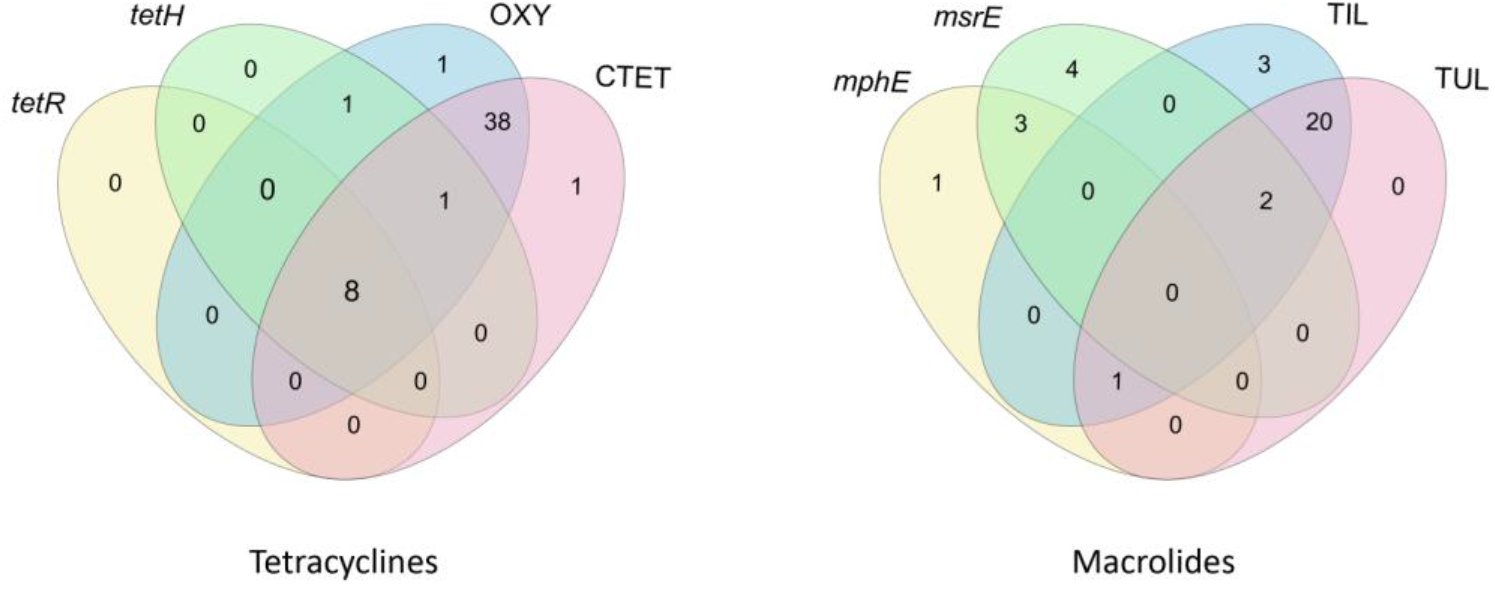
Venn diagrams showing the cooccurrence of antimicrobial resistance genes and phenotypic resistance in the studied strains of *P. multocida* and *M. haemolytica*. The common parts indicate the types of configurations, while the numbers in the respective fields represent the frequency of each configuration. CTET – chlortetracycline, OXY – oxytetracycline, TUL – tulathromycin, TIL – tilmicosin

## Discussion

Bovine respiratory disease (BRD) is the leading cause of antimicrobial use in cattle farming [21–23]. The increasing resistance to antimicrobials represents a significant concern, not only for the health of calves but also for human health [24–29]. Recent molecular studies conducted by our team have revealed that *P. multocida* and *M. haemolytica* are the most frequently identified pathogens among Polish calves exhibiting symptoms of BRD [10].

The most commonly sold antimicrobials in the EU are penicillins (mainly extended-spectrum penicillins in Poland) and tetracyclines, which together account 56.2% of the aggregated sales for food-producing animals by antimicrobial class in European countries in 2022. Compared with other countries in the European Union, Poland has a relatively high antimicrobial consumption [24].

To the best of the authors’ knowledge, there is a lack of published studies on the antimicrobial susceptibility of *P. multocida* and *M. haemolytica* isolated from Polish calves with BRD. This gap in the literature highlights the urgent need for comprehensive studies to better know the resistance profiles of these pathogens.

We present and discuss antimicrobials for which breakpoints have been established by the CLSI in the context of susceptibility categories (S, I, R). On the other hand, for antimicrobials for which breakpoints are not defined by the CLSI, we discuss the data in terms of MIC_50_ and MIC_90_ values.

### *P. multocida* -serotype detection and virulence-associated genes

The study we conducted on *P. multocida* was designed in accordance with the methodology outlined by Townsend et al. It was intended to detect the specific *kmt1* gene and identify all capsular genotypes.

However, our results revealed the presence of only genotype A. Based on available literature and preliminary (unpublished) studies, we determined that lipopolysaccharide 3 is the most prevalent [40]. This finding was confirmed in our study, as we detected it in 93.8% (45/48) of *P. multocida* isolates. Studies conducted on symptomatic Spanish cattle also revealed that the majority of isolates belonged to serotype A:L3, accounting for 97.6%. of all isolates (166/170) [41]. Similar results from other studies show that A:L3 was commonly detected in *P. multocida* isolates associated with BRD [18, 42–46].

All *P. multocida* strains examined in our study exhibited the presence of the *sodA* gene, which helps bacteria evade the host immune response. In the studies from Spain, Germany and Japan this gene was also detected at a rate of 100% in *P. multocida* serotype A strains [33, 41, 44]. In studies from Iran, this gene was less frequently observed in *P. multocida* strains of other serotypes (B, E), occurring at a rate of 63.6% [47]. Studies from Iran, where other serotypes in addition to A were also present, demonstrated a complete absence of this gene [29].

Studies by Ewers et al. have demonstrated a correlation between the presence of the *hgbB* gene and the manifestation of clinical symptoms in cattle. Moreover, this gene was found in 57.7% (104/180) of isolates from cattle [33]. The *hgbB* gene, which may indicate the bacterium’s ability to utilize host haemoglobin, was found in 37.5% (18/48) of the strains investigated in our study. A high prevalence of this gene, at 74.0% (176/238), was reported by Katsuda et al. among strains from diseased calves [44]. Conversely, in the study by Calderón Bernal et al., the prevalence was 0.6% (1/170) [41].

The *ompH* gene encodes a protein that aids bacteria in adhering to host tissues, thereby facilitating colonization. In our study, it was the least frequently detected virulence gene in *P. multocida*, present in only 20.8% (10/48) of the strains. Similar results to ours were obtained by Calderón Bernal et al., who reported a prevalence of 14.1% (24/170), whereas Ewers et al. and Katsuda et al. reported 100% presence of the gene [33, 41, 44]. Interestingly, Gulaydin determined that if this gene is present alone (without any other VAGs cooccurring), it is not associated with the manifestation of the disease. In our study, however, it was consistently found in conjunction with other VAGs that we investigated.

We did not find a statistically significant correlation between the *ompH* and *hgbB* genes. Similarly, Ewers et al. did not report such a correlation and also found no correlation between the other VAGs included in our study [33].

### *M. haemolytica* -serotype detection and virulence-associated genes

In our study, 86.4% (19/22) of the *M. haemolytica* isolates were genotypically classified as serotype A2, while the remaining 13.6% (3/22) were classified as serotype A1. No serotype A6 was detected, despite conducting the analysis according to the protocol outlined by Klima et al., which included primers for all three genes determining serotypes [35, 37].

*M. haemolytica* serotype A2 was considered nonpathogenic to cattle and was regarded as a commensal of the normal respiratory flora [35, 48]. Current research provides updated insights into the role of this serotype; it is also recognized as an opportunistic pathogen capable of causing disease in calves. It was clearly associated with fatal acute lung disorder in calves in the Netherlands, with all of these *M. haemolytica* isolates (n = 49) belonging to serotype A2 [19]. Moreover, the study by het Lam et al. reported that 96.1% (49/51) of the isolates from symptomatic calves were serotype A2, while all 45 isolates from cows were classified as serotypes A1 and A6 [19].

In the study by Mason et al., *M. haemolytica* serotype A2 was identified in 29.8% (31/104) of the *M. haemolytica* isolates tested, which were obtained from bovine clinical pathology and postmortem samples associated with pneumonia cases in Great Britain [49].

Studies conducted in Denmark by Kudirkiene et al. showed that serotype A1 was detected equally in healthy 52.4% (11/21) and diseased calves 47.6% (10/21), whereas serotype A2 was more frequently isolated from diseased calves 70.0% (7/10). In addition, similarly to our study, they did not detect serotype A6 [18].

Abed et al., reported that 60% (9/15) of *M. haemolytica* isolates were classified as serotype A2, whereas 40% (6/15) were identified as serotype A1; all of these strains were obtained from pneumonic calves presenting with respiratory symptoms [50]. Similarly, in the present study focusing on diseased calves, the majority of *M. haemolytica* isolates were identified as A2, including one sample obtained from lung tissue, whereas serotype A1 was considerably less common in our study.

Interestingly, the study by het Lam et al. demonstrated that two serotype A2 strains were genetically closer to serotypes A1 and A6 than to other serotype A2 strains [19]. Such results underscore the need for further research to elucidate the role of these serotypes in the pathogenesis of BRD in calves.

Furthermore, all the *M. haemolytica* strains examined in our study exhibited the presence of the VAGs investigated. These genes are essential for the pathogenicity of *M. haemolytica*, influencing its ability to cause disease and interact with the host’s immune system. One of the most important virulence factors is considered to be the bacterium’s ability to produce leukotoxin, which has cytotoxic potential and damages host leukocytes [20].

Abed et al. demonstrated the presence of the *lkt* and *gcp* genes in 80% (4/5) of the strains obtained from diseased calves [50]. The *gcp* gene encodes a glycoprotein involved in bacterial adhesion to host tissues, facilitating colonization and infection [20]. In a separate study by Dokmak et al., both the *gcp* and *gs60* genes were detected in all strains from cattle exhibiting symptoms of BRD [51]. The *gs60* gene encodes a heat shock protein that enables the bacterium to withstand hostile environmental conditions, enhancing its survival [20].

### Antimicrobial susceptibility -subsceptible, intermediate, resistant

Tetracyclines, which are classified under Category D in the EMA classification—indicating that they should constitute the first line of treatment—exhibit the highest resistance rates among *P. multocida* strains, with resistance levels ranging from 79.2% to 81.3% [52]. Similarly, among the *M. haemolytica* strains analysed, half of them exhibited nonsusceptibility to tetracyclines. The resistance of these bacterial species to tetracyclines has been reported in numerous studies worldwide. It is quite common for resistance rates to exceed 80%, reaching up to 100%, particularly among *P. multocida* strains [4, 21, 29, 53–56]. Compared to those obtained in our study, lower resistance rates were reported by Dutta et al. (4.2%-40.8%). Interestingly, the abovementioned study also revealed that resistance among the studied bacteria is markedly higher when they originate from the lower respiratory tract [57].

In contrast, the *M. haemolytica* strains presented the highest resistance to PEN and TIL, with rates of 63.6%, whereas *P. multocida* exhibited resistance rates of 18.8% and 25.5% for these antimicrobials, respectively. High levels of resistance to TIL have also been reported globally for both species [21, 53, 55, 58]. However, some studies indicate higher resistance rates for *P. multocida*, ranging from 73.1% to 77.3% [4, 53, 55]. Conversely, other reports on TIL resistance among *M. haemolytica* revealed considerably lower resistance rates, ranging from 0.6% to 36.1% [4, 54, 57].

However, the resistance to PEN among *M. haemolytica* reported in other studies is not as high as that observed in our study; similarly, resistance among *P. multocida* has also been reported at relatively low levels [4, 21, 54, 55]. Notably, in studies of cattle from Denmark, all the examined strains of *P. multocida* and *M. haemolytica* were found to be susceptible to PEN and TIL [18]. Considering nonsusceptibility as a combination of resistant and intermediate strains, it is noteworthy that the highest level of nonsusceptibility recorded in our study was to PEN, with an observed rate of 86.4% among *M. haemolytica* strains. Furthermore, this antimicrobial showed the most statistically significant difference in resistance levels between the studied bacteria.

In our study, the highest susceptibility was observed for fluoroquinolones: *P. multocida* presented a 91.7% (44/48) susceptibility rate to enrofloxacin, while 77.3% (17/22) of *M. haemolytica* strains were susceptible to both ENRO and DANO. Our results differ from those of other studies, which reported high percentages of resistant strains for both bacterial species with respect to the aforementioned fluoroquinolones [4, 21, 29, 53, 54, 56, 58]. However, our findings are consistent with those of Klima et al., who also observed low levels of resistance to fluoroquinolones [55]. It is beneficial that we observe low resistance rates to fluoroquinolones, as ENRO, DANO and SPE, which ranked next in terms of susceptibility, fall under Category B in the EMA classification [52]. This classification signifies that these agents are categorized as ‘restricted’ and emphasizes the ongoing need for careful stewardship in their application.

### MIC_50_ and MIC_90_

For the bacteria we studied, which were sourced from the respiratory tract of cattle, the breakpoint for resistance to AMP was established by the CLSI at 0.25 µg/mL. The concentration range on the plates we used starts precisely at this level. Consequently, it was not possible to assess the susceptibility of the examined strains to this antimicrobial using CLSI guidelines. Nonetheless, for both bacterial species, the MIC_50_ was 0.25 µg/mL, with the MIC_90_ reaching 8 µg/mL (Figure 2). In the studies by Katsuda et al., a narrower disparity between these metrics was observed: the MIC_50_ for *P. multocida* was 1 µg/mL, while the MIC_90_ was 4 µg/mL [44]. In contrast, for *M. haemolytica*, this disparity was more pronounced, with the MIC_50_ at 2 µg/mL and the MIC_90_ reaching as high as 128 µg/mL [59].

Moreover, for both bacterial species in our study, the MIC_50_ for NEO was the same as the lowest concentration tested, i.e., 4 µg/mL. Additionally, for *M. haemolytica*, this concentration also corresponds to the MIC_90_.

In contrast, the situation is different for TYLT and CLI. Most of the tested strains did not exhibit growth inhibition or did so only at the highest concentrations tested. However, this may be attributed to the intrinsic resistance of these bacterial species to these substances.

Elevated MIC values for TYLT and CLI were also reported by Depenbrock et al.; however, their study differs from ours in the concentrations of NEO, where the MIC_50_ and MIC_90_ values were notably higher than those observed in our study, exceeding 32 µg/mL [53]. Similarly, high concentrations of 32 µg/mL and above for NEO and TYLT were reported by Andres-Lasheras et al. [4].

### Multidrug resistance

Another significant issue is the simultaneous occurrence of resistance to various classes of antimicrobial agents. In our study, the resistance to at least one substance in three or more antimicrobial classes was 27.1% (13/48) for *P. multocida*, while among *M. haemolytica* strains, resistance was notably higher at 40.9% (9/22). The resistance rate for *P. multocida* in our study was relatively low compared with that reported in other studies, whereas the resistance level for *M. haemolytica* fell within the moderate range.

Among dairy heifers in California, the MDR rates were 76% (110/145) for *P. multocida* and 70% (83/119) for *M. haemolytica* [53]. In contrast, among Canadian dairy cattle, the MDR rates were 67.8% (164/242) for *P. multocida* and 25.4% (53/209) for *M. haemolytica*[4].

Klima et al. have reported varying levels of MDR ranging from 50% to 90% for *P. multocida* and 55% to 81.8% for *M. haemolytica* across different years [55]. Additionally, the majority of serotype A2 strains from calves with fatal acute lung disorders (95.6%) harbored multiple antimicrobial resistance genes, which contributed to three to five antimicrobial classes, including phenicols, sulphonamides, tetracyclines, aminoglycosides, and beta-lactams [19].

### Configurations of resistance patterns

Klima et al. identified 104 unique multidrug resistance patterns involving 16 different antimicrobials [55]. In our study, we observed 25 configurations, although the number of antimicrobials considered was 11. Similarly, Depenbrock et al. reported the same number of configurations as our study, while examining the same number of antimicrobials.[53].

In our study, the most frequently observed phenotypic resistance pattern was ‘CTET, OXY’. However, this pattern was exclusively observed among *P. multocida* strains, occurring in 37.5% (18/48) of the strains. Furthermore, these are antimicrobials agents belong to the same class.

In contrast, among the *M. haemolytica* strains, resistance to ‘XNL, CTET, OXY, PEN, TIL, TUL’ was the most frequently observed configuration and was present in 18.2% (4/22) of the tested strains. Although this configuration involves six antimicrobials, they belong to four distinct classes. Depenbrock et al. found that *P. multocida* was most commonly resistant to four classes, while *M. haemolytica* was resistant to three classes, which contrasts with our findings [53].

Furthermore, their study revealed that resistance to fluoroquinolones was present in the most common configurations for both *P. multocida* and *M. haemolytica*, which also distinguishes their findings from ours. In contrast, in our study, configurations containing ENRO and/or DANO were observed in only 8.6% (6/70) of the results [53].

They also noted a similarity in the resistance patterns, suggesting that this might be due to both bacterial species being isolated from the nasal cavity [53]. In our study, however, these patterns appeared to differ, and we have demonstrated statistically significant differences in antimicrobial resistance between these two bacterial species.

### Resistance genes

In our study, the presence of the *tetH* and *tetR* genes was consistently associated with phenotypic to tetracycline resistance, but the genes *mphE* and *msrE* were detected not only in strains exhibiting phenotypic resistance to macrolides but also in those that did not show such resistance (Figure 4). Similar findings were reported by Klima et al. [55]. The *tetR* gene was always found in cooccurence with the *tetH* gene in the examined strains (n=8). Each of these strains exhibited phenotypic resistance to the both tested tetracyclines (CTET and OXY). Among all the strains in which the *tetH* gene was detected, phenotypic resistance to OXY was observed in 10 out of 10 cases, one of these strains did not show resistance to CTET (1/10). Cooccured resistance to both CTET and OXY in the presence of the tetH gene was noted in 9 out of 10 strains. These findings are consistent with those of the study by Klima et al., which demonstrated a statistically significant association between the presence of *tetH* and resistance to OXY [55].

In our study, resistance genes for antimicrobials such as *tetR, tetH, mphE*, and *msrE* were relatively rare. Therefore, we were not able to make inferences based on correlation tests. However, phenotypic resistance to tetracyclines and macrolides was observed at a high level (Figure 1). The observed antimicrobial resistance may result from the presence of various genes as well as other factors, such as biofilm production or mutations [60, 61]. More than 40 genes responsible for tetracycline resistance are known; for ex ample among cattle the *tetG* gene was found in Germany, *tetL* in Belgium, *tetB* in France and *tetH* emerged as the most frequently detected gene in North America [55, 61–66].

Understanding the mechanisms underlying resistance in the strains studied would require more extensive research.

### Limitations

This study comprises only clinical samples submitted to a single Polish laboratory, which primarily receives specimens from the southwestern region of the country. Due to financial constraints, we were unable to perform LPS typing for three *P. multocida* strains that were negative in the confirmation test for LPS type three. Additionally, the small number of *M. haemolytica* serotype A1 strains (3/22) precluded the possibility of conducting reliable comparative analyses between serotypes.

### Future directions

In this study, the MIC_90_ values for more than half of the tested antimicrobials fell outside the range of the concentrations examined. Furthermore, the statistically significant differences observed in the susceptibility of the bacterial species to antimicrobial agents complicate the formulation of recommendations for field veterinarians regarding the most appropriate first-line antimicrobials to use for BRD in calves. Despite the high susceptibility of the bacteria to fluoroquinolones, it is important to note that these drugs are highly classified by the EMA and should not be used indiscriminately. Therefore, we believe that microbiological testing, including determining the susceptibility of isolated bacterial strains, should precede any decision to use antimicrobial agents in calves in Poland.

The levels of average antimicrobial resistance among *M. haemolytica* are relatively high, raising some concerns. It is intriguing to consider whether, over the next few years, these values may evolve into full resistance and what factors might contribute to this trend. Such dynamics in resistance could significantly impact treatment efficacy and necessitate the development of new therapeutic strategies. It will be important to monitor these trends in the future to better understand the mechanisms of resistance evolution and to implement appropriate preventive measures.

The potential role of *M. haemolytica* serotype A2 as a pathogen, which was previously considered exclusively nonpathogenic in cattle, makes it an intriguing subject for further research. Investigating its role in respiratory diseases could provide valuable insights to aid in the development of prevention strategies.

## Conclusion

The results of this study expand the knowledge of the pathogenicity and antimicrobial resistance of *P. multocida* and *M. haemolytica* in Polish cattle. One third of the strains showed multidrug resistance. *P. multocida* exhibited the highest resistance to tetracyclines, while *M. haemolytica* demonstrated the greatest lack of sensitivity to penicillin. Both bacterial species were found to be susceptible to fluoroquinolones.

The findings from the current study revealed the multidrug resistance of the isolated strains and the presence of genes related to the virulence and resistance of *P. multocida* and *M. haemolytica* bacteria isolated from calves with respiratory symptoms.

## Decalarations

### Consent for publication

Not applicable.

### Availability of data and materials

The datasets used during the current study are available from the corresponding author on reasonable request.

### Funding

The APC is financed by Wroclaw University of Environmental and Life Sciences.

### Authors’ contributions

Conceptualization, A.L.-W. and K.R.; Data Curation, A.L.-W.; Formal Analysis, A.L.-W.; K.D.; Investigation, A.L-W., A.C., I.P., M.K., M.S; Methodology, A.L.-W., A.C., I.P., M.K., M.S; M.D.K.-B., K.D. and K.R.; Resources, A.L.-W. and K.R.; Supervision, M.D.K.-B. and K.R.; Visualization, A.L.-W.; Writing—Original Draft, A.L.-W.; Validation, M.D.K.-B. and K.R.; All authors have read and agreed to the published version of the manuscript.

